# Single-cell RNAseq reveals the pro-regenerative role of senescent FAPs in muscle regeneration

**DOI:** 10.1101/2023.12.12.571148

**Authors:** Cheng Chen, Marielle Saclier, Jérémy Chantrel, Sébastian Mella, Aurélie Chiche, Han Li

## Abstract

Muscle regeneration is associated with transient induction of cellular senescence. However, the role of senescence in muscle regeneration of young mice remains unclear. Using a mouse model deficient in both Cdkn1a and Cdkn2a, we find that a marked reduction in senescent cells correlates with delayed muscle regeneration. Single-cell RNA sequencing reveals a heterogeneous senescence program composing of multiple cell types. Notably, senescent fibro-adipogenic progenitors (FAPs) upregulate Mcl-1 to acquire apoptosis resistance. Moreover, removing senescent FAPs using a Mcl-1 inhibitor S63845 impairs muscle regeneration. Furthermore, we find that senescent FAPs promotes myogenic differentiation in a paracrine manner. Hence, these results highlight the beneficial role of senescent stromal cells in supporting muscle regeneration.

## BACKGROUND

Cellular senescence is a unique cell state induced by stress, which is collectively characterized by irreversible cell-cycle arrest, a set of phenotypic changes, and acquisition of a dynamic senescence-associated secretory phenotype (SASP) composed of a complex mixture of cytokines, chemokines, extracellular matrix remodelling factors, and growth factors^1^. Permanent growth arrest is a potent cell-intrinsic tumor suppression mechanism comparable to apoptosis, whereas SASP mediates crosstalk with the microenvironment^2^. It has become increasingly clear that senescence phenotypes, particularly *in vivo*, are highly heterogeneous depending on the initial stimuli, affected cell type, and spatiotemporal context^3^. Senescent cells can arise at every stage of life and positively contribute to embryonic development^4,5^, tissue repair^6^, and tumor suppression in a temporary manner. However, persistent accumulation of senescent cells has been suggested to drive aging and many aging-associated diseases^7^.

The skeletal muscle displays a unique regenerative capacity from entirely damaged to fully restored histology and function in less than a month. Thus, this tissue provides one of the most powerful systems to investigate how distinct tissue-resident stem/progenitor cell populations orchestrate both intrinsic and extrinsic regulators during tissue regeneration, aging, and pathology^8^. Notably, muscle stem cells (MuSCs), which are indispensable for muscle regeneration, become senescent during aging, which largely accounts for their functional decline ^9,10^. Importantly, ablation of senescent cells or suppression of senescence induction has been shown to improve muscle regeneration in old mice ^9–13^. Interestingly, muscle injury induces an acute senescence response^14^, and we previously demonstrated that senescence promotes cellular plasticity during reprogramming in muscle ^15,16^. However, the role of senescence in physiological muscle regeneration remains debatable ^11,17, 18^.

Here, we characterized the heterogeneous senescence program and its functional relevance during muscle regeneration. Using the *Cdkn1a^-/-^;Cdkn2a^-/-^*mouse model, we found that blunt senescence induction was associated with delayed muscle regeneration. Through single-cell transcriptomic profiling, we found that fibro-adipogenic progenitors (FAPs) are a major senescent cell type in regenerating muscles. Notably, senescent FAPs upregulate Myeloid Cell Leukemia 1 (Mcl-1) to acquire resistance to apoptosis. Furthermore, the pharmacological inhibition of Mcl-1 eliminates senescent FAPs and delays muscle regeneration. Hence, these findings indicate that senescent stromal cells promote muscle regeneration by creating a pro-regenerative niche to support myogenic functions.

## RESULTS

### Muscle injury induces a transient senescence response

We first determined the dynamics of senescence induction during the muscle regeneration. The *Tibialis Anterior* (TA) muscles from mice of 8-12 weeks old were injured using cardiotoxin (CTX) and collected on different days post-injury (DPI) (Figure 1A). We co-stained the muscle sections with SAβGal and F4/80, a pan-macrophage marker, to distinguish macrophages from other cell types. SAβGal signal was first detected at 3DPI, peak at 5DPI, and became non-detectable after 14DPI (Figure 1B). Of note, most SAβGal+ cells were positive for F4/80 at 3DPI and 5DPI, while SAβGal^+^/F4/80^-^ cells became more apparent at 10DPI (Figure 1B). Next, we used C_12_FDG, a fluorogenic substrate for β-galactosidase ^19^, in combination with the macrophage marker CD64 to quantify and isolate cells via fluorescence-activated cell sorting (FACS) (Figure 1C, Supplementary Figure 1A). Consistent with immunohistochemical staining, a non-macrophage C_12_FDG^+^ population emerged at 5DPI and persisted till 10DPI (Figure 1D and 1D’).

**Figure 1.**
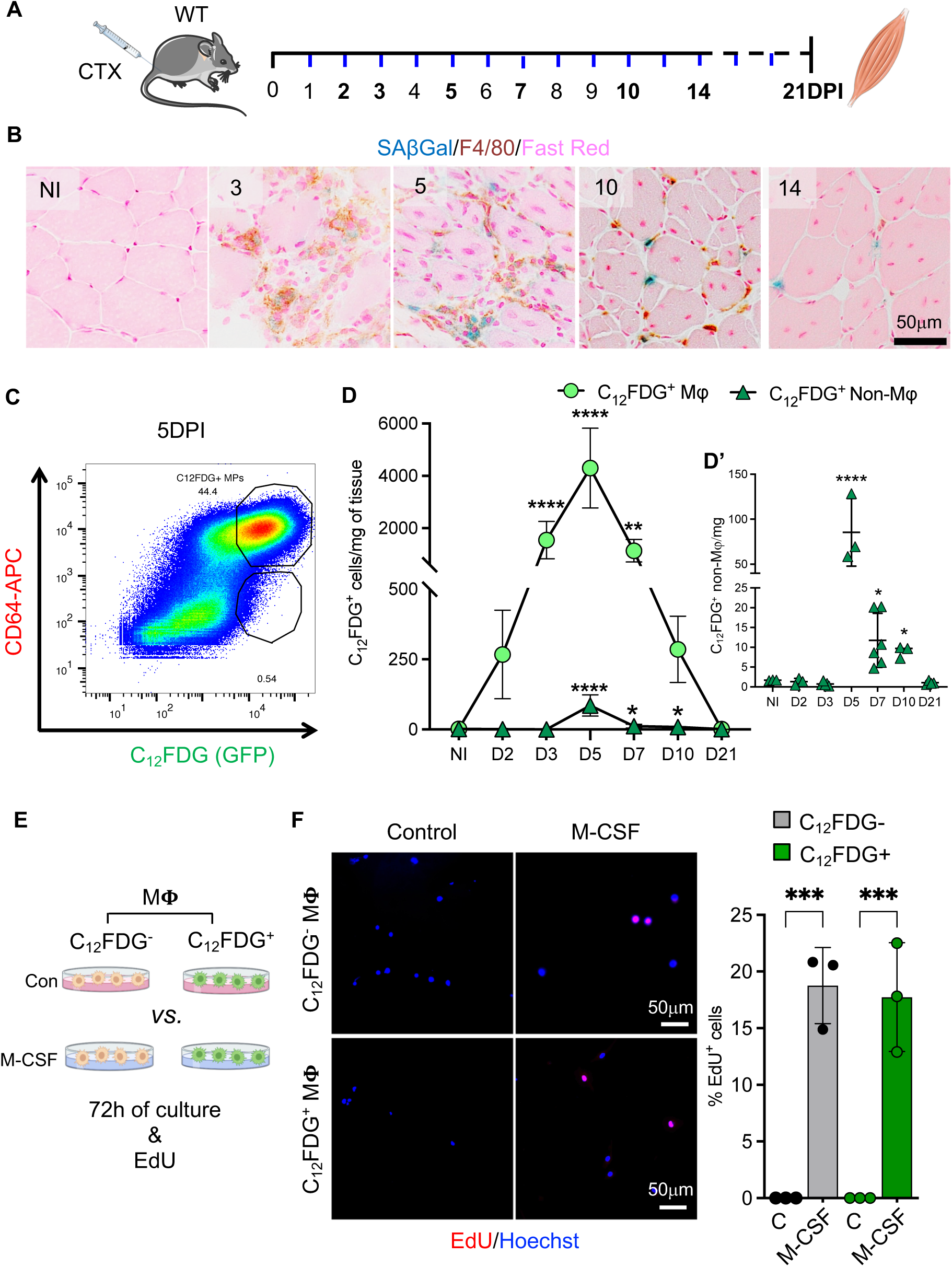
Senescence and macrophages during muscle regeneration. **A.** Experimental scheme for determining senescence in regenerating muscle. **B.** Representative images of SA-β-Gal & F4/80 co-staining in TA cross-sections at multiple time points. Scale bar=50μm. **C.** Representative FACS plot of cells derived from 5DPI and stained with CD64-APC & C12FDG. **D & D’.** Numbers of C12FDG+ macrophages (Mϕ) and non-macrophages per mg of tissue at different time point during muscle regeneration. n=3-5 mice (2 TAs of the same mouse pooled for 1n). **E.** Experimental scheme of *in vitro* culture of macrophage. **F.** Representative images and quantification of EdU incorporation by C12FDG+ or C12FDG-Mϕ after 72h with or without M-CSF. n=3 Scale bar=50μm; For D, data are means ± SD, for F, data are means ± SD of at least three independent experiments. Statistical significance was determined using a two-tailed Student’s t-test for both D/D’ and F. ****p < 0.0001, ***p < 0.001. See also **Figure S1**.

Given that macrophages exhibit increased lysosomal activity independent of senescence ^20^, we determined whether C_12_FDG+ macrophages are senescent in our system. Firstly, most C_12_FDG+ macrophages at 5DPI were Ly6C^low^, a marker associated with the pro-regenerative phenotype (Supplementary Figure 1C and 1D). Secondly, we observed an increase mRNA expression of Glb1 and Cdkn1a, but a reduction mRNA expression of Cdkn2a, in C12FDG+ vs. C12FDG-macrophage (Supplementary Figure 1E). Thirdly, loss of LaminB1 expression is a marker associated with senescence ^21^, and there was a significant reduction of LaminB1 intensity in the non-macrophage C_12_FDG+ cells, while no difference was observed within macrophage subpopulations regardless of C_12_FDG intensity (Supplementary Figure 1F and 1G). Finally, we isolated both C_12_FDG+ and C_12_FDG-macrophages and cultured them with M-CSF to stimulate survival and proliferation (Figure 1F). Interestingly, there was a comparable percentage of EdU+ cells in both populations, indicating that C_12_FDG+ macrophages retained their proliferative capacity, at least *in vitro*, upon mitogenic stimulation (Figure 1E and 1F). Taken together, our data suggest that C12FDG positivity might reflect a feature of macrophages independent of senescence, consistent with the report that elevated β-galactosidase activity is associated with polarized macrophage ^22^. Therefore, in the current study, we focused on the non-macrophage populations.

### Delayed muscle regeneration correlates with senescence reduction in Cdkn1a^-/-^; Cdkn2a^-/-^ mice

Next, we explored the functional impact of senescence during muscle regeneration using mouse models lacking p16 and/or p21 (Supplementary Figure 2A). p21 and p16 can act redundantly in mediating senescence induction in certain *in vivo* contexts, such as wound healing ^6^ and tumour suppression ^23^. Therefore, we compared the senescence levels in the regenerating muscles of *Cdkn1a^-/-^*, *Cdkn2a^-/-^*, and *Cdkn1a^-/-^/Cdkn2a^-/-^* (DKO) mice with WT. Interestingly, we found a significant reduction in SAβGal+ cells in *Cdkn1a^-/-^* and DKO mice, but not in *Cdkn2a^-/-^* mice, compared to WT at 10DPI (Figure 2A). In addition, DKO exhibited a consistent reduction of SAβGal+ cells at earlier time points of 5 & 7DPI (Supplementary Figure 2B). Consistently, analysis of whole muscle extracts revealed decreased expression of senescence-associated mRNAs such as *Pai-1, Plau,* and *Tnfaip6* in DKO mice (Supplementary Figure 2C). Notably, there was a slight yet significant reduction in p57 expression, but no difference in p27, and a marked increase in p15 expression (Supplementary Figure 2C), indicating potential compensatory mechanisms with other cyclin-dependent kinase inhibitors (CKIs) in DKO.

**Figure 2.**
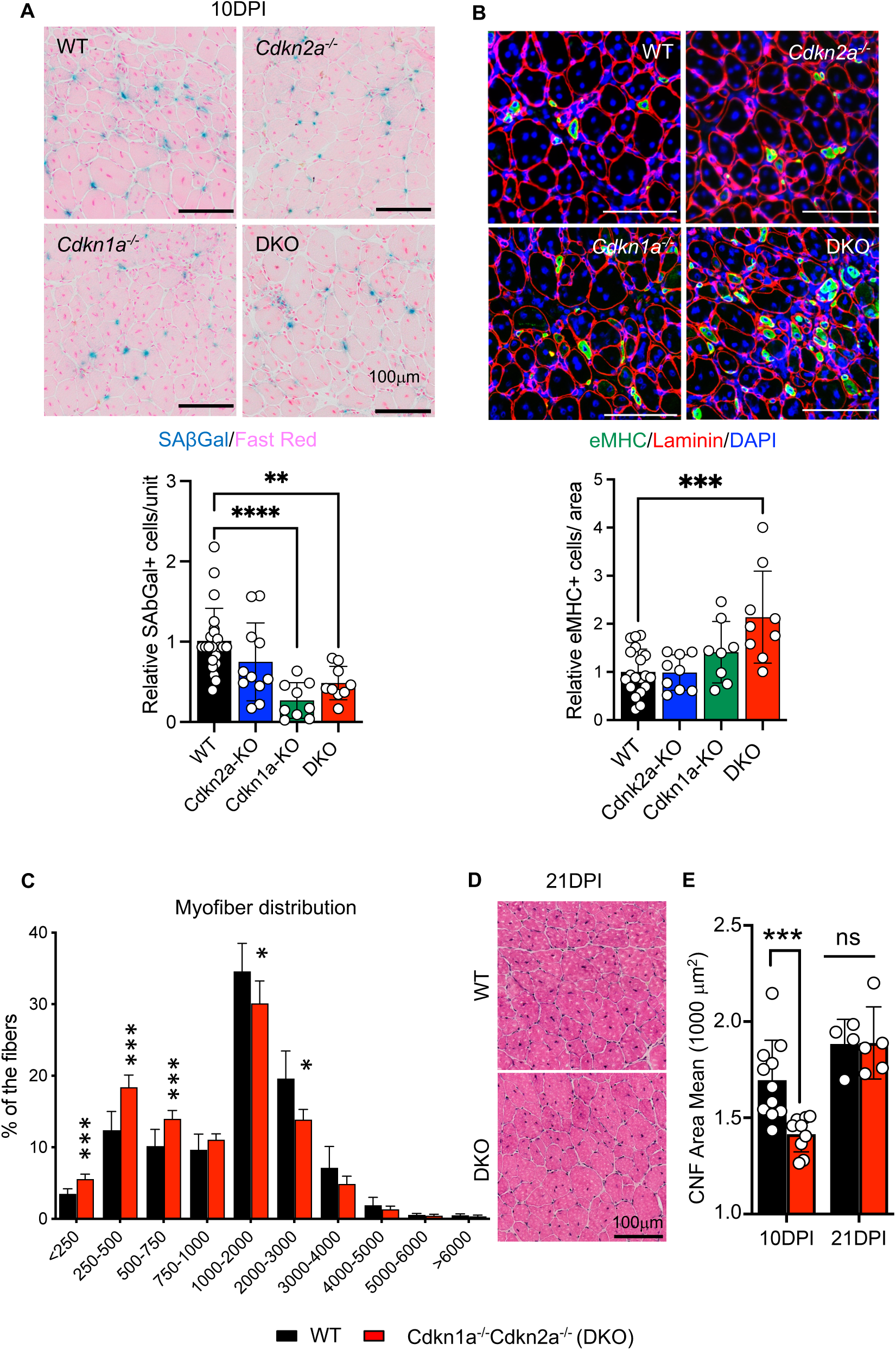
Reduced senescence and delayed regeneration in Cdkn1a/Cdkn2a DKO. Representative images and quantification of SAβGal. **A.** Laminin and embryonic myosin heavy chain (eMyHC) stainings. **B.** staining of TA muscle cross-section from mice of indicated genotypes at 10 DPI. *Lower panels*, quantification of SAβGal+ cells (**A.**) and eMyHC+ cells. **B.** normalized to WT. n=10 mice for every genotype, one value one TA/mouse. Scale bar=100μm. **C.** Quantification of CSA and frequency distribution analysis of the centrally nucleated fibres (CNF) in TA muscle of WT and DKO at 10DPI, 1TA/mouse. **D.** Representative images of haematoxylin and eosin (H&E) staining of TA muscles from WT and DKO at 21DPI, a time point when muscle regeneration is consider completed. Scale bar=100μm. **E.** Quantification of mean CSA of central nucleated fibres between WT and DKO of 10DPI and 21DPI. TAs from 11 mice for WT and 9 mice for DKO at 10DPI, 4 mice for WT and 5 mice for DKO at 21DPI. 1TA/mouse. Scale bar,100μm. Error bars represent mean ± SD. Statistical significance was determined using Mann-Whitney test, *p <0.05, **p<0.001, *** p<0.0001.

We next evaluated the muscle regeneration capacity of mutant mice. Embryonic myosin heavy chain (eMyHC) is a specific marker of immature fibers that is exclusively expressed in newly regenerating myofibers, indicating early muscle regeneration. At 10DPI, WT samples contained a limited number of eMyHC+ fibers. Interestingly, there was more eMyHC+ fibers in the DKO compare to the other samples (Figure 2B). Besides, we observed a trend of increased eMyHC+ fibers in the Cdkn1a^-/-^ but not in the Cdkn2a^-/-^ samples (Figure 2B). Of note, delayed muscle regeneration has been reported in the Cdkn1a^-/-^ mice previously ^24^. In addition, myofiber growth, measured by the cross-sectional area (CSA), was greatly retarded in the DKO compared to WT (Figure 2C & 2D). Moreover, there was increased expression of the immature fiber marker, Myh8, and decreased expression of the mature fiber marker, MyoZ1, in DKO muscle extracts compared to WT (Supplementary Figure 2D). Notably, there was no difference in the number of Pax7+ MuSCs (Supplementary Figure 2E) between WT and DKO at 10DPI. Finally, there was no difference in fiber size between DKO and WT at 21DPI, when the muscle was fully regenerated (Figure 2D & 2E). Taken together, we concluded that muscle regeneration was delayed in DKO mice compared to WT mice, highlighting a correlation between reduced senescence and delayed muscle regeneration in mice lacking both Cdkn1a and Cdkn2a loci.

### Transcriptomic analysis of the senescence program during muscle regeneration

To determine the causal role of senescence in muscle regeneration and better understand the senescence program *in vivo*, we characterize the senescence program during muscle regeneration using both bulk RNA sequencing (RNA-seq) and single-cell RNA sequencing (scRNA-seq) (Figure 3A). We focused on 10DPI, a later time point during muscle regeneration, based on two considerations: 1/SAβGal+ cells are readily detectable at 10DPI; 2/ minimized non-senescence-specific staining of C12FDG. For bulk RNA-seq, we compared the transcriptomes of two populations, C12FDG+ & F4/80-(S2) *vs.* C12FDG-(S1). Principal component analysis (PCA) revealed a clear separation between S1 and S2, indicating distinct biological properties (Supplementary Figure 3A). Volcano plot analysis identified 1164 genes with significant differential expression (Supplementary Figure 3B). Gene-set enrichment analysis (GSEA), using a recently established gene set that identifies senescent cells across tissues (SenMayo_Mouse)^25^, showed significant enrichment of senescence-associated genes in S2 compared to S1 (Figure 3B). A heatmap was generated containing a collection of senescence-associated genes that were found to be significantly upregulated in S2 (Figure 3C). In addition, GSEA using the hallmark gene sets revealed that several pathways that are frequently associated with senescence, including coagulation, protein secretion, and inflammatory response, were significantly enriched in S2 compared to S1 (Supplementary Figure 3B).

**Figure 3.**
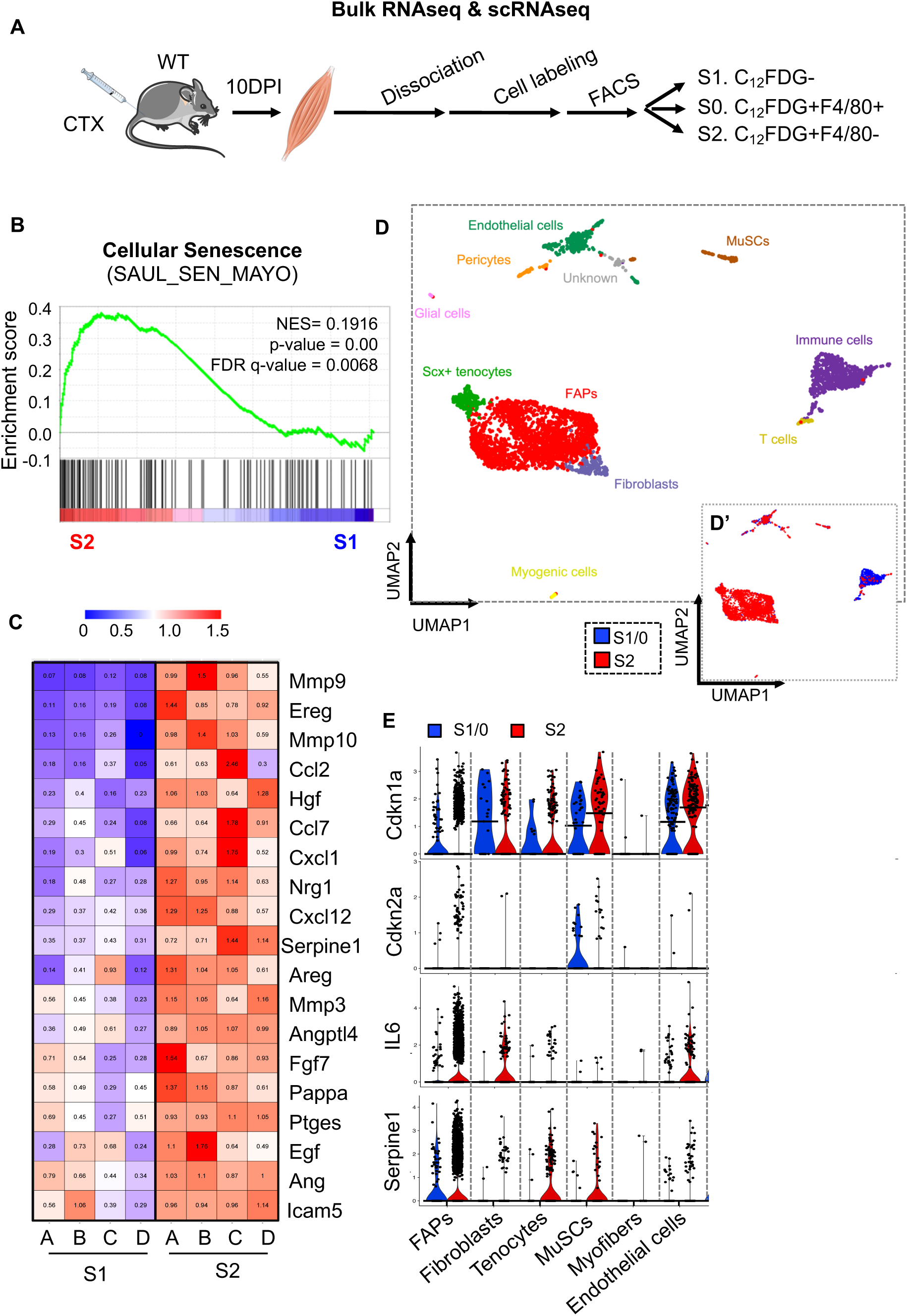
Transcriptomic analysis of the senescence program. **A.** Scheme of the experiments. **B.** GSEA enrichment plot of the cellular senescence signature in S2 (C12FDG+) vs. S1 (C12FDG-). **C.** Heat map of enriched senescence-related genes. **D.** Uniform Manifold and Projection (UMAP) plot of the mononuclear cell populations identified by scRNA-seq in regenerating muscle of 10DPI (data summary consisted of n=3 young male mice). **D’.** Visualization of the abundance between C12FDG+ and C12FDG-populations. **E.** Expression of *Cdnk1a*, *Cdkn2a*, *Il-6*, and *Serpine1*(Pai1) across indicated cell populations.

To determine the senescent cells’ transcriptional heterogeneity during muscle regeneration, we performed scRNA-seq on two populations: pooled C12FDG-(S1) and C12FDG+; F4/80+ (S0) populations (refereeing as new S1), and C12FDG+; F4/80-(S2), to enrich non-macrophage C12FDG+ cells while retaining the C12FDG+; F4/80+ population. After quality control and data filtration, we obtained 1077 cells from S1 and 3174 cells from S2 for subsequent analysis. Eleven clusters were identified by cluster analysis and visualized using the uniform manifold approximation and project (UMAP; Figure 3D). All the cell types described in the regenerating muscle were detected (Supplementary Figure 3D)^26^. We also identified sub-cluster 9, which was positive for several proliferation markers. However, owing to the low cell number, we were unable to determine its identity. We checked the distribution of S1 and S2 populations across cell types. The S2 population contained most of the clusters, except T (cluster 8) cells, and macrophages were exclusively detected in S1 due to arbitrary redistribution (Supplementary Figure 3E). Notably, FAPs, tenocytes, and fibroblasts were more enriched in S2 than in S1 (> 10 folds, Figure 3D’, Supplementary Figure 3E). Next, we examined the expression of p16 and p21 in the scRNA-seq dataset. Consistent with a previous report ^13^, we found that p16 was only detected in very few FAPs and muscle stem cells (MuSCs), whereas p21 was evident in most cell types (Figure 3E). Notably, we observed a substantial increase in the number of cells expressing higher levels of p21 in S2 than in S1 in multiple cell types, including FAPs, fibroblasts, tenocytes, MuSCs, and endothelial cells (Figure 3E), which correlated with the elevated expression levels of other senescence-associated genes, such as IL-6 and Pai1 (Figure 3E). Taken together, both bulk RNA-seq and scRNA-seq datasets indicate that multiple cell types acquire senescence-like features during muscle regeneration.

### FAPs is a major senescent cell type in regenerating muscle

To understand the direct impact of senescence on muscle regeneration, we focused on FAPs, which are the dominant cell type in S2 (Supplementary Figure 3E). scRNA-seq analysis indicated that FAPs from S2 upregulated the expression of various senescence-associated genes compared with FAPs from S1 (Supplementary Figure 4A). To validate the scRNA-seq results, we examined various senescence markers using freshly isolated FAPs from PDGFRa-H2B::eGFP reporter mice ^27^. Given that the number of FAPs peaks around 3-4DPI^28^ and the C_12_FDG+/F4/80-population peaks around 5DPI (Figure 1D), we isolated FAPs from TAs of both non-injured (NI) and 5DPI. After 24 hours of culturing *in vitro*, FAPs were stained with several markers (Figure 4A). Compared to the NI control, the 5DPI FAPs exhibited an increase in SAβGal+ (Figure 4B) and γH2AX+ cells (Supplementary Figure 4D), and a decrease in LaminB1+ and BrdU+ cells (Supplementary Figure 4B & 4C). Therefore, FAPs acquire senescence in a DNA damage-dependent manner in regenerating muscle.

**Figure 4.**
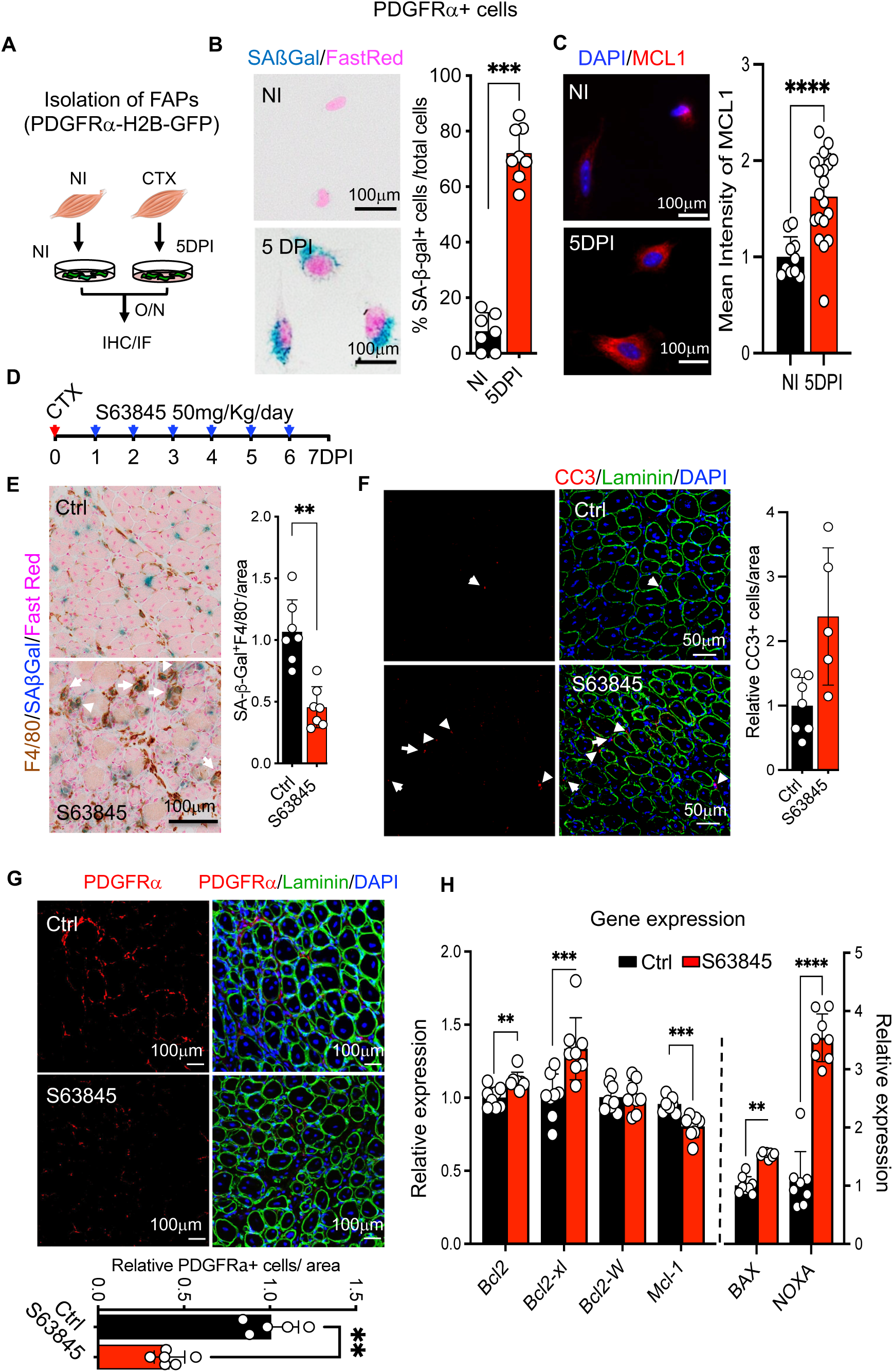
S63845 induces apoptosis in senescent FAPs *in vivo*. **A.** Experimental scheme. **B & C.** Representative images and quantifications of indicated staining of isolated FAPs from WT TA muscle of non-injured and 5DPI. B. SAβGal, n=3; **C.** Mcl-1, n=3. Scale bar=100μm **D.** Experimental scheme. **E-G.** Representative images and quantifications of indicated staining of control vs. S63845 treated TA muscles at 7DPI. **E.** SAβGal/F4/80 co-staining, n=7; **F.** Cleaved Caspase 3 (CC3)/Laminin co-staining, n=7; **G.** PDGFRα/Laminin co-staining, n=5. Scale bar=100μm **H.** Quantitative PCR analysis of the expression of indicated genes. Error bars represent mean ± SD. Statistical significance was determined using Mann-Whitney test, *p <0.05, **p<0.001, *** p<0.0001.

### Mcl-1 inhibitor eliminates senescent FAPs during muscle regeneration

Next, we sought to eliminate senescent FAPs by using small molecules to directly evaluate their impact on muscle regeneration. As senescent cells can acquire apoptosis resistance via upregulation of Bcl-2 anti-apoptotic family members ^29^, we compared the gene expression of Bcl2, Bcl2xl, BclW, and Mcl-1 in the FAPs cluster. Interestingly, we observed a marked increase in Mcl-1 expression but not in Bcl2, BclxL, and BclW (Supplementary Figure 4E). Importantly, we found that FAPs from 5DPI expressed higher levels of Mcl-1 protein compared to NI (Figure 4C). It has recently been reported that Mcl-1 is an alternative target for killing senescent cells ^30,31^. Therefore, we explored whether inhibition of Mcl-1 could eliminate senescent FAPs during muscle regeneration. We treated regenerating muscle with S63845, a specific Mcl-1 inhibitor, for six days and harvested the TAs at 7DPI (Figure 4D). We found a significant reduction in SAβGal+/F4/80-cells in S63845 treated TAs compared to the controls (Figure 4E). Importantly, there were more apoptotic cells (Figure 4F) and fewer FAPs in the treated TAs group than in the controls (Figure 4G). Notably, numerous F4/80+ macrophages surrounded the SAβGal+ cells upon S63845 treatment (Figure 4E), suggesting a robust response to apoptotic cells. Analysis of whole muscle extracts revealed upregulation of the pro-apoptotic genes BAX and NOXA, further confirming the elevated apoptosis in S63845-treated samples (Figure 4H). Curiously, we observed a decrease in Mcl-1 and an increase in Bcl2 and Bcl2-xL expression, suggesting a potential compensation among Bcl-2 anti-apoptotic family members (Figure 4H). Collectively, inhibition of Mcl-1 induces apoptosis in senescent FAPs during muscle regeneration.

### Reduction of senescent FAPs impairs muscle regeneration

We next evaluated the impact of eliminating senescent FAPs on muscle regeneration. Histological examination revealed that S63845-treated TA muscles contained persistent inflammatory infiltration, necrotic myofibers, and cellular debris, which were largely absent from control samples at 7DPI (Figure 5A). S63845-treated TA muscles exhibited widespread eMyHC expression (Figure 5B), retarded myofiber growth (Figure 5C and Supplementary Figure 5A), and altered gene expression profiles (Figure 5D). At the cellular level, we observed a significant reduction in endothelial cells, whereas no difference was observed in Pax7+ MuSCs in S63845-treated TA muscles compared to the control (Supplementary Figure 5B & 5C). Collectively, these results demonstrated that eliminating senescent FAPs delays muscle regeneration.

**Figure 5.**
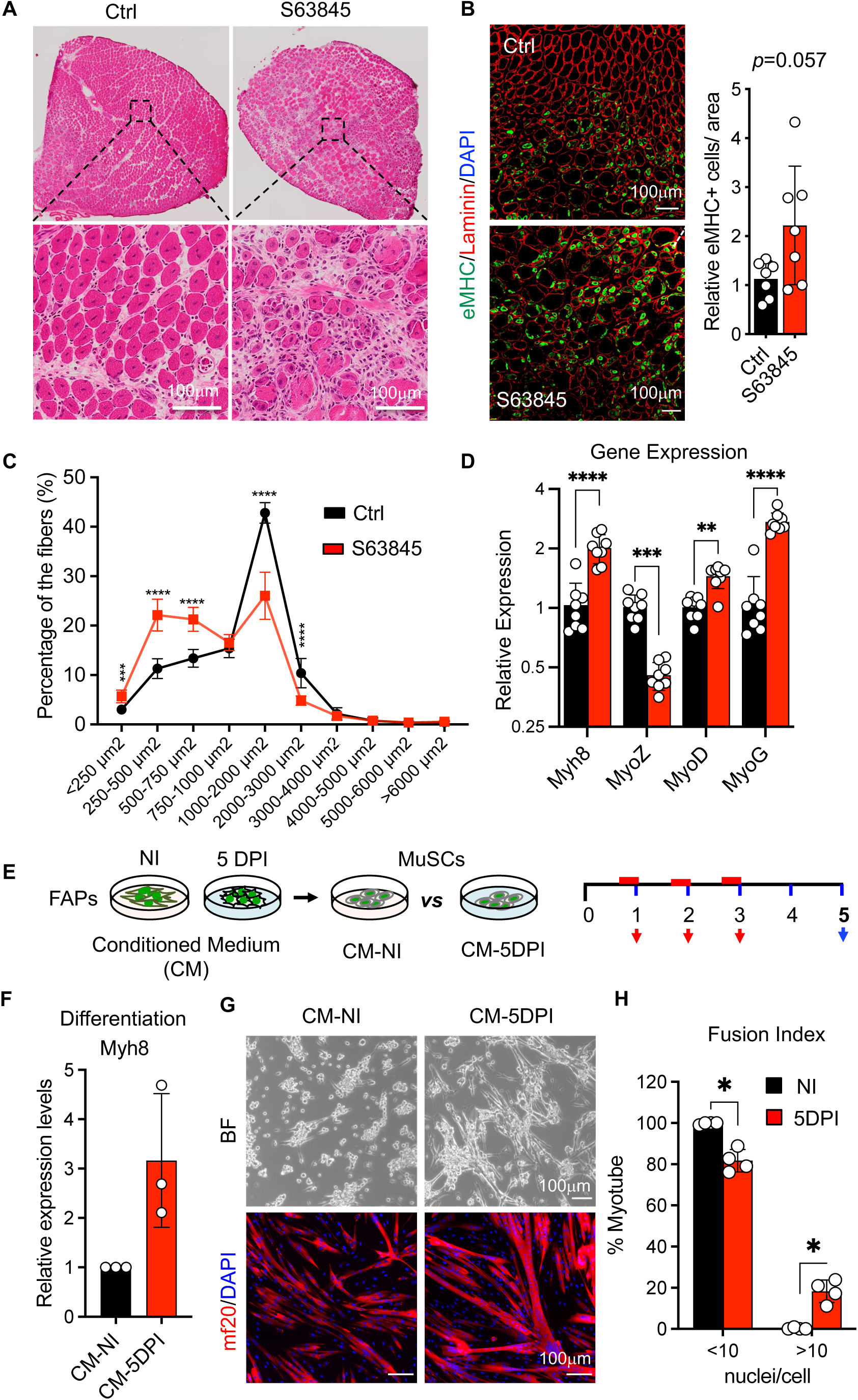
S63845 delays muscle regeneration. **A.** Representative images of H&E staining of TA muscle from the control group and the S63845 treatment group at 7DPI. Scale bar=100μm **B.** Representative images and quantification of eMyHC and Laminin co-staining in TA muscle. n=7, 1TA/mouse from control and S63845 treated, Scale bar=100μm. **C.** Quantification of CSA and frequency distribution analysis of the centrally nucleated fibres (CNF) in TA muscle of control and S63845 treated at 7DPI, n=7, 1 TA/mouse. **D.** Expression of indicated genes. **E.** Experimental scheme. **F.** Quantitative PCR analysis of Myh8 gene expression. 6-8 mice/experiment, n=4. **G.** Bright field (upper panel) and MF20 immunofluorescence staining (lower panel) of myoblasts cultured with FAPs CM derived from TAs of either NI or 5DPI. Scale bar=100μm **H.** Quantification of fusion capacity by fusion index evaluation using MF20/nuclei stained images of G. 6-8 mice/experiment, n=4. Error bars represent mean ± SD. Statistical significance was determined using Mann-Whitney test, *p <0.05, **p<0.001, *** p<0.0001.

### Senescent FAPs promote MuSCs differentiation *in vitro*

FAPs secrete signaling molecules to support myogenesis ^32,33^, and we have previously shown that senescent cells promote MuSCs plasticity during *in vivo* reprogramming in a paracrine manner ^15^. Therefore, we wondered whether senescent FAPs could facilitate myogenic potential via SASP. To address this hypothesis, we used a well-established *in vitro* culture system in which MuSCs are activated into myoblasts in a proliferation medium and differentiated and fused into myotubes in a differentiation medium. Both proliferation and differentiation media were reconstituted with conditioned medium (CM) of FAPs isolated from either NI or 5DPI muscles to assess the paracrine impact of senescent FAPs on myogenic properties (Figure 5E). EdU incorporation analysis showed no difference between senescent and non-senescent FAPs-CMs, indicating that senescent FAPs did not alter MuSCs activation and proliferation (Supplementary Figure 5D). Interestingly, there were many more myotubes when MuSCs were cultured in differentiation medium containing senescent FAPs-CM (Figure 5F-H), suggesting an enhanced capacity to differentiate and fuse. Taken together, senescent FAPs could facilitate myogenic differentiation and fusion potential via SASP to ensure optimal muscle regeneration.

## Discussion

Here, we show that a reduced senescence response correlates with delayed muscle regeneration in mice deficient for important senescence mediators, p16 and p21. scRNA-seq analysis revealed heterogeneous senescent cell populations, mainly composed of stromal cells, including FAPs, tenocytes, and endothelial cells. We found that senescent FAPs depend on upregulated Mcl-1 for survival, and are important for optimal muscle regeneration, at least in part, by supporting myogenic differentiation.

It has recently been reported that macrophages can acquire senescence phenotypes and promote early lung tumorigenesis ^34,35^. Indeed, we also observed that the majority of C12FDG+ cells upon muscle injury were macrophages (Figure 1C & 1D). However, C12FDG+ macrophages retained their proliferative capacity comparable to that of C12FDG-macrophages *in vitro* (Figure 1F) and did not exhibit several common markers of senescence (Supplementary Figure 1E-1G). Therefore, in the context of muscle regeneration in young mice, increased β-galactosidase activity might reflect an intrinsic property of macrophages, such as polarization, rather than senescence ^20^.

It has been reported that acute ablation of p16+ cells upon muscle injury by treating p16-3MR mouse model with ganciclovir (GCV) resulted in a significant reduction in senescence and improved muscle regeneration ^11^. Surprisingly, we did not observe a significant difference in SAβGal+ cells or muscle regeneration dynamics in *Cdkn2a^-/-^* mice compared to those in WT mice (Figure 2A & 2B). This discrepancy could be explained by the known compensation effect of germline knockout of Cdkn2a ^23^, which yields much weaker phenotypes than acute disruption of gene function ^6^. Of note, we observed a strong reduction in SAβGal+ cells in the regenerating muscles of *Cdkn1a^-/-^* and *Cdkn1a^-/-^; Cdkn2a^-/-^* mice (Figure 2A).

To determine the role of senescence in muscle regeneration, we first found that muscle regeneration was delayed in DKO mice compared to that in WT mice (Figure 2). Second, the direct elimination of senescent FAPs, the major senescent cell type revealed by scRNA-seq (Figure 2D and Supplementary Figure 2E), using a Mcl-1 inhibitor, delayed muscle regeneration (Figure 4E-4G, Figure 5A-C). Third, CM derived from senescent FAPs promoted myogenic differentiation and fusion without altering MuSC activation and proliferation (Figure 5F & 5H, Supplementary Figure 5D), which agrees with the timing of senescence induction upon injury (Figure 1B & 1D). The discrepancy between Moiseeva et al. and our results could potentially be explained by the heterogeneity of the *in vivo* senescence program. It is known that susceptibility to certain senolytic is cell type-dependent ^36^. However, it is currently unknown which cell populations are being eliminated. We found that both FAPs and endothelial cells were markedly reduced following Mcl-1 inhibitor treatment (Figure 4G & Supplementary Figure 5B). However, we do not know whether dasatinib and quercetin, used by Moiseeva et al., target the same cell types. Moreover, neither study was able to completely eradicate senescent cells during muscle regeneration. Therefore, different senescent cell types may play distinct roles in muscle regeneration. Future investigations are warranted to elucidate senescence heterogeneity during muscle regeneration.

Collectively, we characterized the senescence program during muscle regeneration and found that senescent stromal cells, particularly FAPs, promoted optimal muscle regeneration. These findings will further our understanding of senescence heterogeneity and its functional relevance to tissue regeneration.

## ACKNOWLEDGEMENTS

We are indebted to Jun Zhang for his excellent technical support in the image analysis. We are grateful to the Central Animal Facility, Cytometry Platform, and Bioinformatics and Biostatistics Hub of Institut Pasteur. This work was funded by Institut Pasteur, Centre National pour la Recherche Scientific and the Agence Nationale de la Recherche (Laboratoire d’Excellence Revive, Investissement d’Avenir; ANR-10-LABX-73; ANR-16-CE13-0017, ANR-21-CE13-0006-01). Work in the HL laboratory was also funded by Agence Nationale de la Recherche (ANR-22-CE16-0015-03), Foundation ARC (PJA 20181208231), and AFMTELETHON (22403). Marielle Saclier is recipient of Revive postdoc fellowship, Jeremy Chantrel is recipient of DIM Longévité et vieillissement PhD fellowship.

## AUTHOR CONTRIBUTIONS

C.C, ans M.S performed most of the experimental work, contributed to experimental design, data analysis, and discussions. J.C made critical experimental contributions. J.C and S.M performed bioinformatic analysis. AC contributed experimentally. H.L. supervised the study, designed the experimental plan, interpreted the data, and wrote the manuscript. All authors discussed the results and commented on the manuscript. The authors declare no competing financial interests with this paper.

## Supplemental Experimental Procedures

### Mice and breeding

Animals were handled as per European Community guidelines and the ethics committee of the Institut Pasteur (CETEA) approved protocols. *Cdkn2a^-/-^* mice were graciously provided by Manuel Serrano from the Spanish National Cancer Research Centre in Madrid, Spain^1^. The *Cdkn1a^-/-^*mice were obtained from Jackson Lab (JAX: 016565). The Cdkn1a^-/-^; Cdkn2a^-/-^ DKO mice were generated in-house. Pdgfra-H2B-GFP (JAX:007669) and Tg15: Pax7nGFP (JAX:036759) mice were obtained from Institut Pasteur.

### Animal procedures

To induce muscle injury, mice were anesthetized with isoflurane. The tibialis anterior (TA) muscles were injured by injecting 50 μl of Cardiotoxin (CTX), specifically Naja pallida at a concentration of 10 μM (Latoxan, L8102). Following the surgery, mice were administered analgesic buprenorphine (Vetergesic) at a dosage of 0.3 mg/kg. For S63845 treatment, mice were intraperitoneally injected (i.p.) with 50 mg/kg of S63845 (Medchemtronica, HY-100741), or received the vehicle (2% Kolliphor in PBS) (Sigma-Aldrich, C5135)^2^. The treatment was initiated on the same day as the CTX injection and lasted for 6 days. The tibialis anterior muscles were collected on the 7th day after injury (7DPI). All the experiments were conducted with male mice aged between 10 to 14 weeks, unless otherwise specified.

### FACS and Cytometry

#### Isolation senescent cells from regenerating muscle

To isolate senescent cells, injured TA were finely chopped in cold HamF10 (Sigma-Aldrich, N6908) digested in HamF10, 10% FBS medium containing 1,5 U/ml collagenase A (Roche, 11088793001), 3U/ml dispase II (Roche, 4942078001) and 10 ug/ml DNAse I (Roche, 11284932001) for 1h at 37°C under agitation. After incubation, the supernatant was filtered through a 40μm cell strainer (BD Falcon), and red blood cells were lysed using Ammonium Chloride Solution (Stemcell Technologies, 07800). Cells were then incubated 1h at 37°C and 5% CO_2_ in 100nM of bafilomycin A1 (Tebu, 21910-2060). 33μM of C_12_FDG (Fisher scientific, 11590276) was then added for 2h at 37°C and 5% CO_2._ Then, the cells were washed in HamF10, 10% FBS and stained with F4/80-PE (ThermoFischer Scientific, MA516624), CD64-APC (BD Biosciences, 558539), or Ly6C-APC (Fischer Scientific, 12-5931-82) at 4°C for 30 minutes. After additional washes, the cells were resuspended in PBS, 2% FBS before sorting or analysing, Propidium Iodide (Sigma-Aldrich, P4864) was used to discriminate live cells from dead cells. For scRNAseq, senescent cells (C_12_FDG^+^/ F4/80^-^ cells), macrophages (C_12_FDG^+^/F4/80^+^ and C_12_FDG^-^/F4/80^+^ cells) and non-senescent cells (C_12_FDG^-^/ F4/80^-^ cells) were sorted with a 100 μm nozzle using ASTRIOS Sorter. In some experiments, C_12_FDG^-^/CD64^-^, C_12_FDG^+^/CD64^-^, C_12_FDG^-^/CD64^+^ and C_12_FDG^+^/CD64^+^ sorted cells were cytospined 15min at 2000 rpm on Starfrost slides and immunostained (see bellowed). In some experiments, C_12_FDG^-^/CD64^-^, C_12_FDG^+^/ CD64^-^, C_12_FDG^-^/CD64^+^ and C_12_FDG^+^/CD64^+^ sorted cells were centrifugated 15min at 2000 rpm, supernatant was removed and 1ml of Trizol (Fischer Scientific, 12044977) was added for RNA extraction. For cellular populations analysis, C_12_FDG^-^/CD64^-^, C_12_FDG^+^/CD64^-^, C_12_FDG^+^/CD64^+^ and C_12_FDG^-^/CD64^+^ cells were isolated at different time points after cardiotoxin injury were analysed on a LSR Fortessa, BD. In some experiments, Ly6C^+/-^ cells were analysed within the C_12_FDG^+^/CD64^+^ and C_12_FDG^-^/CD64^+^ cell populations.

### Cell culture conditions

#### Isolating satellite cells (SCs) and non-injured FAPs

Isolation of satellite cells (SCs) from Tg15: Pax7nGFP/B6N mice and non-injured fibro/adipogenic progenitors (FAPs) from Pdgfra-H2B-GFP mice was conducted following a previously described protocol^3,4^. Briefly, the muscles from the whole leg were finely chopped in cold HamF10 medium supplemented with 10% horse serum. The chopped muscles were then transferred into a 50-ml tube containing 40 ml of freshly prepared Digestion Buffer 1, which was incubated at 37°C for 45 minutes. Digestion Buffer 1 consisted of HamF10 medium (Sigma-Aldrich, N6908), 10% Horse serum (GIBCO16050122), and 800 U/ml Collagenase CLS2 (Worthington, LS004176). After washing with HamF10 medium, the tissue was incubated in Digestion Buffer 2, containing 100 U/ml Collagenase CLS2 and 1.1 U/ml Dispase II (Fisher Scientific 17105041), for 30 minutes. The digestion was performed by gently pushing the tissue up and down 7-10 times using a 20-ml syringe with a 20-gauge needle. The digested tissue was then filtered through a 40-μm cell strainer. Subsequently, the cells were incubated in Ammonium Chloride Solution (Stemcell Technology 7800) for 1 minute to lyse red blood cells, followed by a wash with HamF10 medium containing 2% horse serum. The cells were now ready for sorting. Propidium Iodide (Sigma-Aldrich, P4864) was used to discriminate live cells from dead cells.

#### Isolating FAPs from Injured TA

The 6-10 TAs were isolated from Pdgfra-H2B-GFP mice at 5 DPI. The protocol was modified according to the previous publication.^5^ Briefly, the TAs were finely chopped in cold HamF10 medium supplemented with 10% horse serum. The chopped muscles were then transferred into a 50-ml tube containing 30 ml of freshly prepared Digestion Buffer, which was incubated at 37°C for 60 minutes. Digestion Buffer consisted of HamF10 medium (Sigma-Aldrich, N6908), 10% Horse serum (GIBCO16050122), and 2mg/ml Collagenase A (Roche11088793001), 3u/ml Dispase II (Sigma-Aldrich, 4942078001) and 10ug/ml DNAase (Roche 10104159001). The digested tissue was then filtered through a 40-μm cell strainer. Subsequently, the cells were incubated in Ammonium Chloride Solution (Stemcell Technology 7800) for 1 minute to lyse red blood cells, followed by a wash with HamF10 medium containing 2% horse serum before sorting. Propidium Iodide (Sigma-Aldrich, P4864) was used to discriminate live cells from dead cells.

SCs and FAPs were collected after sorting and directly cultured in the following medium: DMEM (GIBCO31966), 20% FBS (GIBCO, 10270), and 1% Penicillin-Streptomycin (GIBCO15140). FAPs were seeded on gelatin-treated plates at a density of 5×10^4^ cells/well in a 6-well plate. The FAPs conditioned medium (CM) was collected daily for three days after 24 hours, filtered through 0,2μm filter, stored at −20 °C as described previously^6^. For SCs, the plates were treated with Matrigel (1mg/ml) for 15 minutes and incubated at 37°C for 1 hour before seeding. SCs were seeded at a density of 3×10^4^ cells per well in a 6-well plate, directly with the FAPs CM supplemented with 2% Ultroser G (SATORIUS, 15950-017). From 4^th^ day to 6^th^ day, SCs were cultured with the differentiation medium with DMEM and 2% Horse serum. The cells were fixed at 6^th^ day for further staining. For macrophage proliferation assay, C_12_FDG^+^/CD64^+^ and C_12_FDG^-^/CD64^+^ cells were sorted and cultured in DMEM Glutamax (Fischer Scientific, 11594446), 20% FBS (GIBCO, 10270), 1% Penicillin-Streptomycin (GIBCO15140) with or without 30% of L929 cell line-derived conditioned medium for 24hour.

### Bulk and single cell RNA-Sequencing

For total RNA sequencing, cells were pellet and RNA extraction was performed right after sort with Qiagen RNA extraction RNeasy Mini Kit. 400ng of 3 replicates/ conditions (Non-senescent and senescent) with a total of 6 samples were sent to BGI company for RNA sequencing with a concentration of 20ng/uL. 40M reads were sequenced/samples. Single-cell RNA-sequencing libraries were prepared using the Chromium Single Cell 3’ reagent kit v2 (10x Genomics) following the manufacturer’s protocol. The cells were pelleted and washed twice with PBS following sorting. Manual cell counting was performed to assess cell morphology. For each of the two conditions, 10,000 cells were loaded onto the 10x Chromium Controller. After library preparation and quantification, the libraries were sequenced on the Illumina NexSeq 500 platform.

### Bioinformatic analysis

For Bulk RNAseq, after filtering the raw reads by removing adaptor sequences, contamination, and low-quality reads, we obtained the FASTQ raw data. Indexes were created using the reference annotation gtf file (Mus_musculus.GRCm39.108.gtf) and the reference genome (GRCm39.dna.primary_assembly.fa) fasta file. To process the raw data, we used RSEM (ver. 1.3.2) which applies Bowtie2 for read mapping. The edgR R package was used to study differentially expressed genes, with the Inv0 group acting as the control. A gene with a false discovery rate (FDR) cut-off < 0.05 and a log2 fold change ≥ 1was considered to be differentially expressed. A principal component analysis (PCA) was performed on the edgR-normalized count data after regularized log transformation to estimate and visualize the variations between samples.

Gene Set Enrichment Analysis (GSEA) was performed to identify enriched pathways and gene sets between different sample groups. According to the SeneQuest website, we generated a new set of genes related to cellular senescence. The mouse hallmark (https://www.gsea-msigdb.org/gsea/index.jsp) and cellular senescence gene sets were used as references for GSEA analysis. The significance of enrichment was assessed using the false discovery rate (FDR) threshold of <0.25 and the nominal p-value threshold of <0.05. To visualize the differentially expressed genes in cellular senescence gene set, heatmaps were generated, where rows represent individual genes and columns represent different sample groups. To visualize the enriched signature pathways, a bidirectional histogram was used, where each bar represents a gene set or pathway, and the height of the bar corresponds to the normalized enrichment score (NES) value.

The scRNA-seq data analysis was conducted using the Seurat package (version 4.3.0) in the R programming environment (version 4.2.2). Initially, the raw sequencing data underwent preprocessing steps, including the exclusion of low-quality cells (<5 genes/cell), genes with low expression levels (<1000 UMIs), and cells with excessive mitochondrial RNA content (>20%) within each dataset (S1 and S2). The remaining cells underwent mRNA count normalization using the ‘NormalizeData’ function with the ‘LogNormalize’ method. Subsequently, the ‘FindVariableGenes’ function with the ‘variance stabilizing transformation (vst)’ method was employed to identify the top 2000 most variable genes in the dataset. To remove undesired technical variation, the dataset was scaled and centered using the ‘ScaleData’ function. Principal component analysis (PCA) was then performed on the scaled data to reduce dimensionality, and the top 30 principal components were selected for downstream analysis. For two-dimensional visualization, non-linear dimensionality reduction was performed using the Uniform manifold approximation and projection (UMAP) technique. Integration of both datasets was conducted using the Harmony method^7^,as a batch effect was observed between the S1 and S2 populations. Cell clustering was performed using the graph-based clustering algorithm implemented in Seurat’s ‘FindClusters’ function, with low resolution (0.15) set to identify distinct cell populations. To annotate the different cell types present in the dataset, the ‘FindAllMarkers’ function was utilized to identify the top 12 expressed gene markers specific to each cluster. Cell types were determined based on the established functions of these marker genes. Cell-type annotation was further confirmed by manual inspection of key marker genes specific for each cell type known to be present in the repairing muscle. Differential gene expression analysis was performed using the ‘FindMarkers’ function to identify genes that were significantly upregulated or downregulated between S1 and S2 conditions in the different cell-type clusters (FAPs, endothelial cells, tenocytes, fibroblasts, and MuSCs).

### Immunohistochemistry and immunofluorescence

TA muscles were isolated from mice and immediately frozen in liquid nitrogen by immersing them in cooled isopentane (Sigma-Aldrich, 320404) for 40 seconds. All samples were then stored at −80°C or directly cryosectioned into 10-µm sections.^8^

For histology, sections were air dried and fixed with 4% PFA in PBS and washed, then stained with Haematoxylin and Eosin (H&E). For immunofluorescence, cells were fixed in 4% PFA for 15 minutes and stained with LaminB1(ab16048, 1:200), Mcl1 (ab32087, 1:100) and mf20 (DSHB, 1:30) after blocking with 3% BSA. For tissue sections, eMyHC (DSHB F1.652, 1:50), PDGFRα (AF162,1:200), F4/80 (MA516624, 1:100), and Cleaved Caspase 3 (Cell Signaling 9661,1:300) were stained without prior fixation. Cells and tissue sections were permeabilized with 0.5% Triton X-100 for 5 minutes. The blocking buffer used contained either 10% Normal Goat Serum (Fisher Scientific, 10189722), 10% Normal Donkey Serum (Sigma-Aldrich, D9663), or 3% BSA. Samples were incubated with primary antibodies overnight at 4°C and subsequently incubated with the corresponding fluorescence-conjugated secondary antibodies for 1 hour at room temperature. After staining with DAPI (1 μg/μL in PBS), the samples were mounted with IMMU-MOUNT (Fisher Scientific, 10662815). Same procedure excepting blocking step was realized on cytospined cells. Images were acquired using an Olympus IX83 microscope and quantified using ImageJ software. LaminB1 intensity was calculated with Correlated Total Cell Fluorescence (CTCF = Integrated Density – (Area of Selected Cell x Mean Fluorescence of Background readings)).

### BrdU and EdU immunofluorescence staining

FAPs were cultured on coverslips coated with Gelatin (Sigma-Aldrich, G1890) in 24-well plates for 24 hours. Bromodeoxyuridine (BrdU) was then added and allowed to incorporate into the cells for 2 hours at 37°C. Following incubation, the cells were washed with Dulbecco’s Modified Eagle Medium (DMEM) and phosphate-buffered saline (PBS). Subsequently, the cells were fixed in cold 70% ethanol at 4°C for 20 minutes to preserve their structure and in 2N hydrochloric acid (HCl) at room temperature (RT) for 20 minutes to denature the DNA. The cells were then washed three times with PBS and blocked in a solution of PBS containing 0.1% Triton X-100, 10% goat serum, and 1% bovine serum albumin (BSA) for 1 hour. Subsequently, the cells were incubated overnight at 4°C with a BrdU antibody (Abcam, ab6326, 1:200 dilution).

Satellite cells were seeded on coverslips coated with Matrigel (Corning, 354234) in 24-well plates for analysis. The cells were collected at 24, 48, and 72 hours after seeding. Edu was incorporated for 3 hours before collection. EdU staining was performed using the Click-iT Edu Imaging Kits (Invitrogen C10339). Briefly, the cells were incubated with EdU (10μM) for 3 hours before fixation for satellite cells, and 4 hours for macrophages. After washing with PBS, the cells were fixed in 4% PFA for 15 minutes. Subsequently, the cells were washed with PBS containing 3% BSA and permeabilized with 0.5% Triton in PBS for 20 minutes. Following another wash with PBS containing 3% BSA, the cells were incubated in the Click-iT reaction cocktail for 30 minutes and then in Hoechst for 30 minutes. Finally, the coverslips were mounted with SL PREP IMMU-MOUNT (Fisher Scientific, 10662815). Images were acquired using an Olympus IX83 microscope and quantified using ImageJ software.

### SAβGal & F4/80 co-staining and quantification

The assay was conducted as described previously^8,9^. Briefly, sections were fixed at room temperature for 10 minutes in PBS containing 2% paraformaldehyde and 0.2% glutaraldehyde. The sections were washed with PBS and incubated in the X-gal solution containing 4 mM K3Fe(CN)6, 4 mM K4Fe(CN)6, 2 mM MgCl2, 0.02% NP-40 (Sigma-Aldrich), and 1 mg/ml X-gal (EUROMEDEX, EU0012) in PBS with a pH of 5.5 at 37°C for 48 hours. Subsequently, the samples were washed with PBS and post-fixed in 4% PFA in PBS for 15 minutes. After additional washes, the samples were blocked in a solution of 10% normal goat serum and 0.2% BSA in PBST (0.3% Triton) and incubated overnight at 4°C with anti-F4/80 antibody (Fisher Scientific, MA516624) at a dilution of 1:100. The secondary antibodies conjugated with peroxidase from Dako (K4003) were incubated with the samples for 30 minutes. Liquid DAB+ (Agilent Technologies, K3468) was used for color development. Finally, the sections were counterstained with Fast-red (VECTOR, H3403) and mounted in Eukitt (Sigma-Aldrich, 03989). The sections were scanned using an OLYMPUS VS120 microscope, and SAβgal-positive cells were quantified using a self-developed program called “showblue”^9^.

### Quantitative real-time PCR

Total RNA was extracted from cells and tissue samples with Trizol (Fisher Scientific, 12044977) following provider’s recommendations. Quantitative real-time PCR was performed using a LightCycler 480 (Roche) using SYBR Green Master (Roche, 04887352001). All values were obtained at least in duplicate, and in a total of at least two independent assays. Calculation for the values was made using the ΔΔCt method, as previously described (Yuan et al., 2006).

### Quantification and statistical analysis

The number of independent experimental replications is reported in the figure legends (n, mean ± sem or mean ± sd). Statistical analyses were performed using Graphpad Prism v9 software. Quantification for laminin and centralized nuclei were performed with Muscle J^10^. Statistical significance was assessed by two-tailed student’s t test. P-value <0.05 was considered as statistically significant.

### Data Resources

The accession number for the proteomic data reported in this manuscript is PXD028590 and the datasets are available at the ProteomeXchange via the PRIDE database and will be accessible upon the acceptance of the manuscript and in Tables S1 of this manuscript.

---

**Figure S1.**
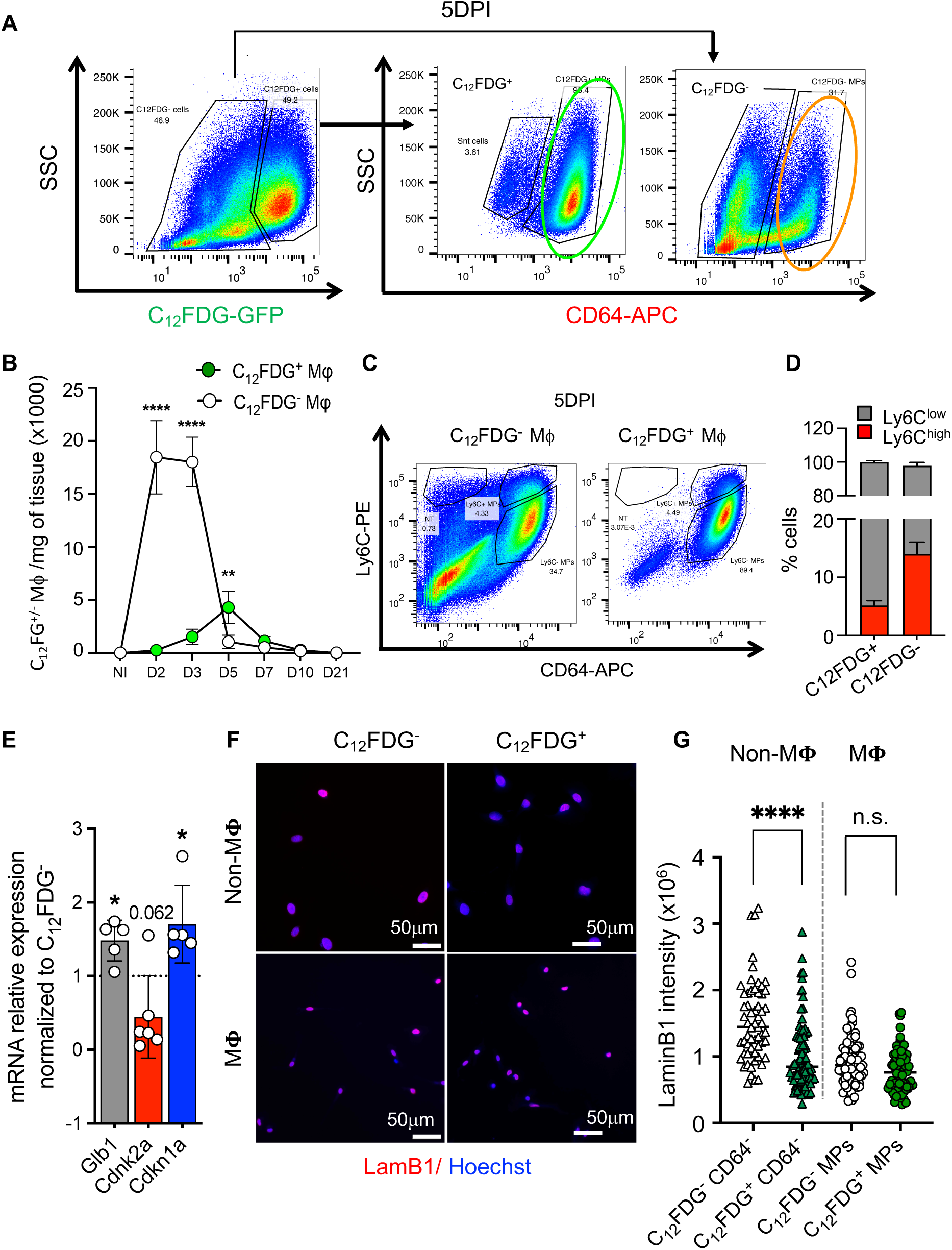
Related to Figure 1. Senescence and macrophages during muscle regeneration. **A.** Representative FACS plots at 5DPI of the gating strategy used to identify C_12_FDG^+^ and C_12_FDG^-^ Mϕ (CD64^+^ cells). **B.** Quantification of number of C_12_FDG^+^ and C_12_FDG^-^ Mϕ per mg of tissue in NI TA and at different time point after muscle injury. n=3-5 mice/time point. **C.** Representative FACS plots of cells stained with CD64-APC and Ly6C-PE form TA at 5DPI. **D.** Percentage of Ly6C^low^ and Ly6C^high^ cells within the C_12_FDG^+^ and C_12_FDG^-^ Mϕ. **E.** Gene expression analysis of indicated genes comparing C12FDG+ to C12FDG-Mϕ. **F.** Representative LaminB1 immunostaining of C_12_FDG^+^ or C_12_FDG^-^ Mϕ and non-Mϕ sorted from TA at 5DPI. Scale bar=50μm. **G.** Quantification of LaminB1 intensity of C_12_FDG^+^ or C_12_FDG^-^ Mϕ and non-Mϕ sorted from TA at 5DPI. Statistical significance was determined using a two-way ANOVA tests for (B) (C_12_FDG^+^ Mϕ vs C_12_FDG^-^ Mϕ), and two-tailed Student’s *t*-test for (E) (C_12_FDG^+^ Mϕ vs C_12_FDG^-^ Mϕ, and C_12_FDG^+^ Mϕ vs C_12_FDG^-^ non-Mϕ). Data are represented as means±SD of at least three independent experiments. * p<0.05, ** p<0,01, **** p<0.0001, n.s. not significant.

**Figure S2.**
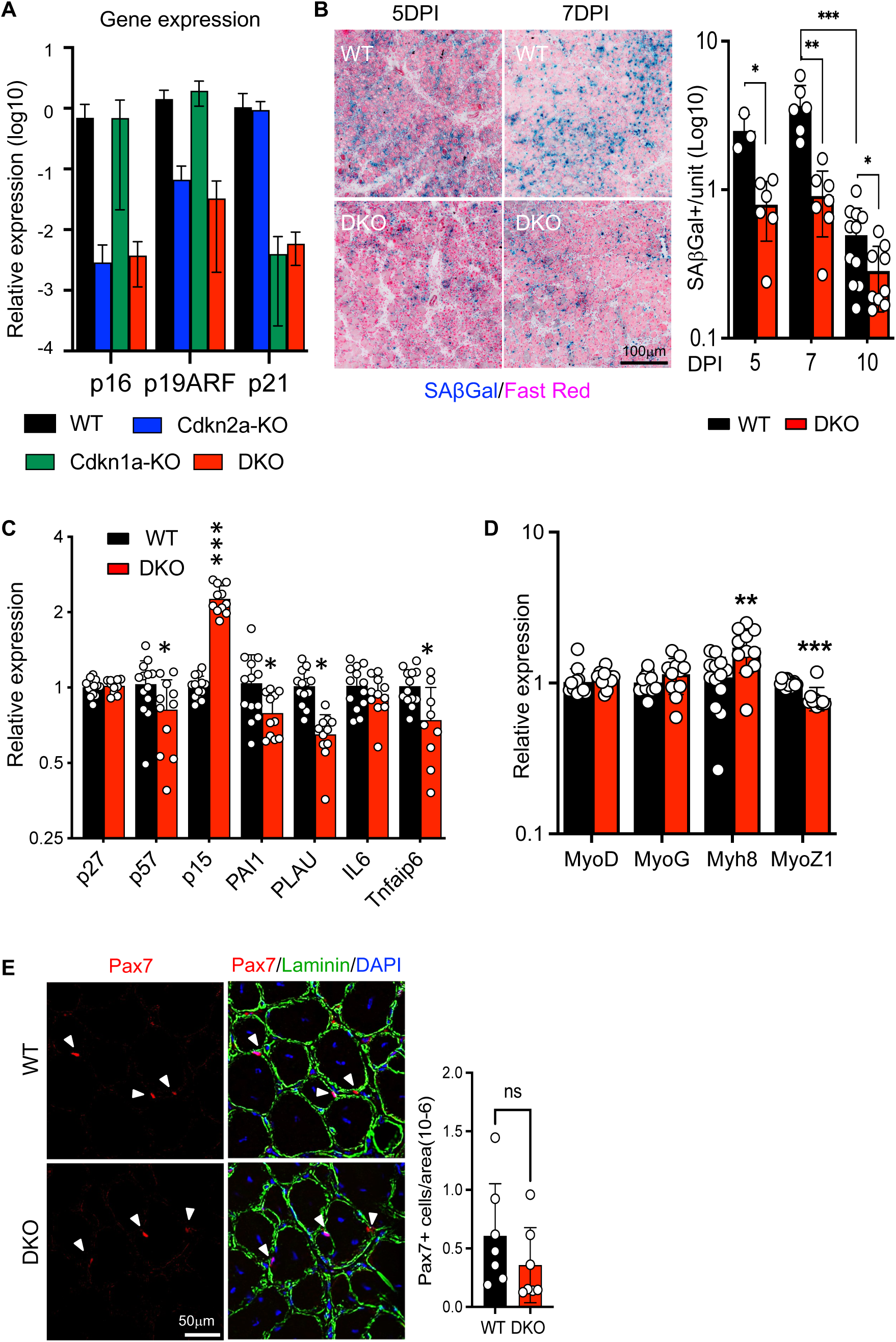
Reduced senescence and delayed regeneration in Cdkn1a/Cdkn2a DKO. **A.** qPCR validation of mouse models. **B.** Representative SAβGal staining of TA muscle cross-section from WT and DKO mice at 5 and 7 DPI, Scale bar=100μm. *Right panel*: quantification of SAβGal+ cells between WT and DKO at different time points during muscle regeneration. n=3-6 mice at 5 DPI. n=6-7 mice at 7 DPI. n=10-13 mice at 10 DPI. **C. D.** qPCR analysis of relative mRNA of indicated genes in TA muscle from WT and DKO at 10 DPI. Senescence-associated (C) and muscle regeneration-associated (D). n= 13 from WT and n=11 from DKO. **E.** Representative images and quantification of Pax7 staining of TA muscle cross-section from WT and DKO mice at 10 DPI. Scale bar=50μm. Statistical significance was determined using Mann-Whitney test. Error bars represent mean ± SD. *p <0.05, **p<0.01, *** p<0.001.

**Figure S3.**
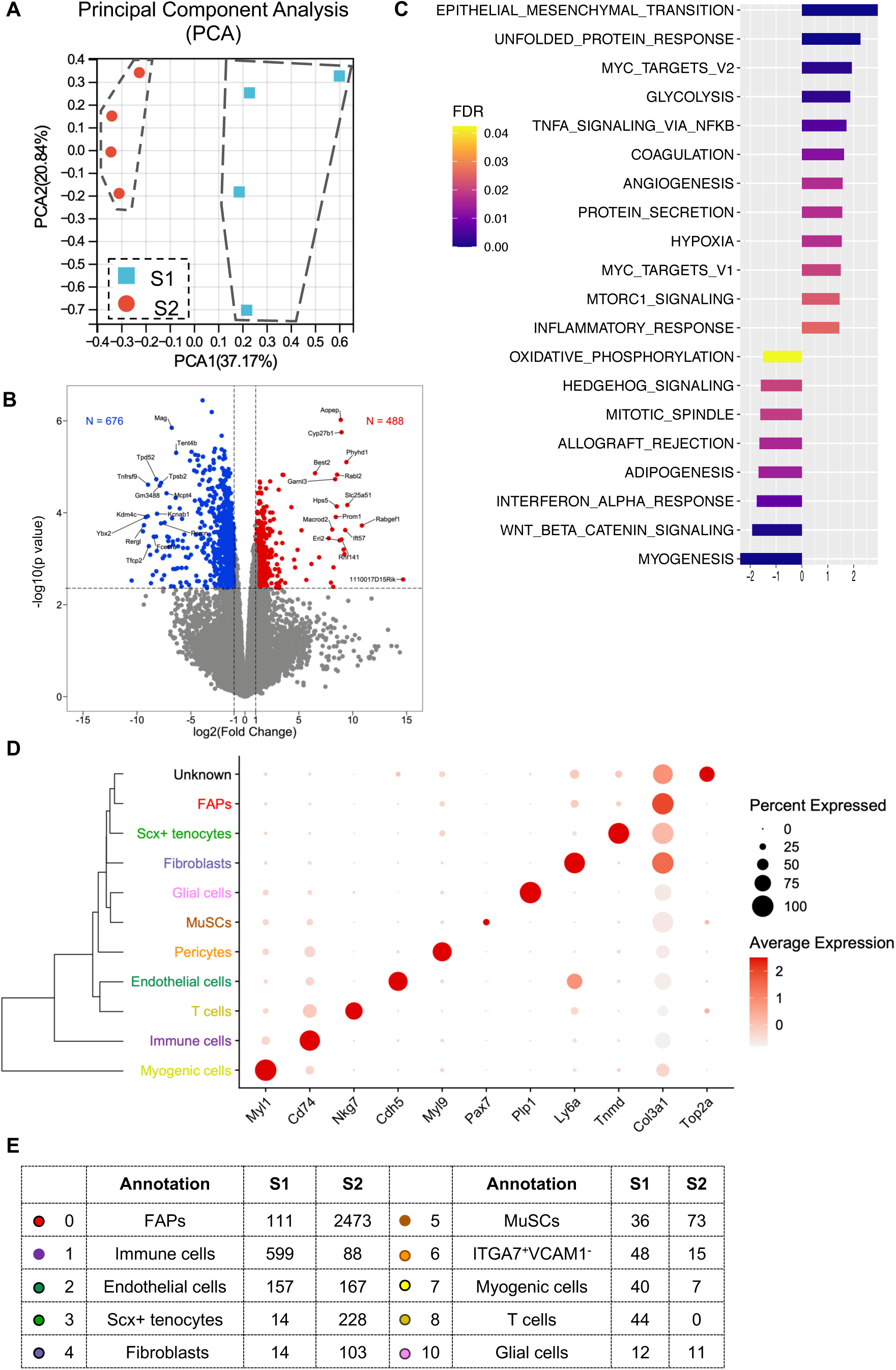
Transcriptomic analysis of the senescence program. **A.** Principal component analysis graph of bulk RNAseq samples. Red circles represent S2 samples, while blue squares represent S1 samples. **B.** Volcano plot illustrating the differential gene expression profile between C12FDG+ non-Mϕ and C12FDG-non-Mϕ cells. Differentially expressed genes are indicated by colored points above the significance threshold (False Discovery Rate < 0.05). Upregulated genes in C12FDG+ non-Mϕ cells are shown in red, while downregulated genes are in blue. Visible gene names are the top 15 upregulated and the top 15 downregulated after ranking using the log2FC x −log10(p-value) formula. **C.** GSEA bar plot of S2 vs S1 differentially expressed genes in FAPs (scRNA seq dataset) using the HALLMARKgene sets collection. **D.** Dot plot illustrating cell type annotation for the scRNA-seq dataset. Genes on the x-axis are examples of key markers for each identified cell types. Dot size represents the percentage of cells expressing marker genes within each cluster, and the color intensity represents the average expression. **E.** Number of cells detected in different cell types between S1 and S2 populations.

**Figure S4.**
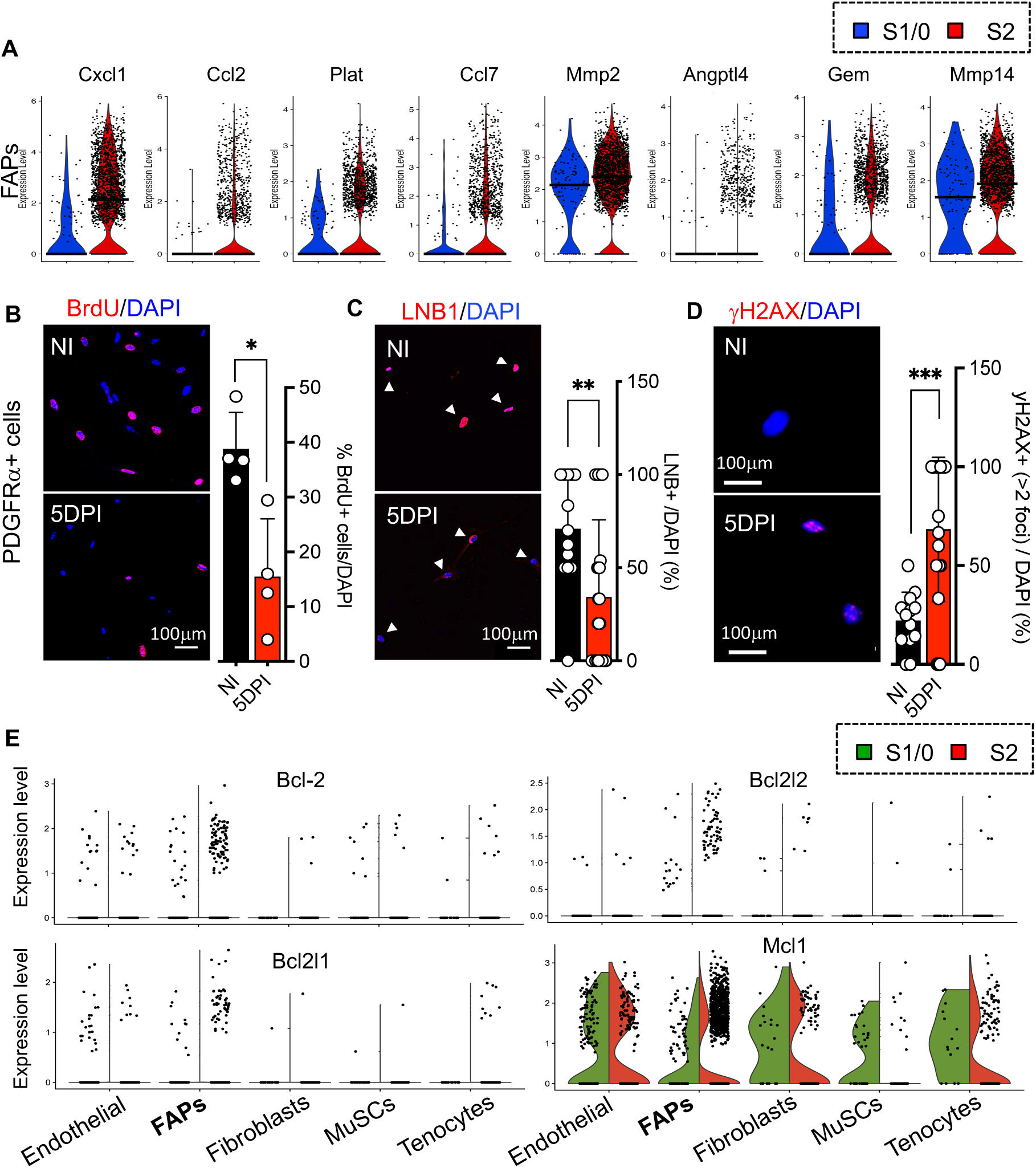
FAPs acquire senescence phenotypes during muscle regeneration. **A.** Violin plots representing the expression of several SASP-related genes in FAPs. **B-D.** Representative images and quantification of BrdU (B), LaminB1(C), and **γ**H2AX (D) stainings in FAPs obtained from both NI and CTX 5 DPI muscle. Scale bar=100μm, the dots represent picture views. **E.** Half violin plots representing the differential expression of Bcl-2 anti-apoptosis family members, Bcl-2, Bcl2l1, Bcl2l2, and Mcl1, in different indicated cell types between S1 and S2. Statistical significance was determined using Mann-Whitney test. Error bars represent mean ± SD. *p <0.05, **p<0.01, *** p<0.001

**Figure S5.**
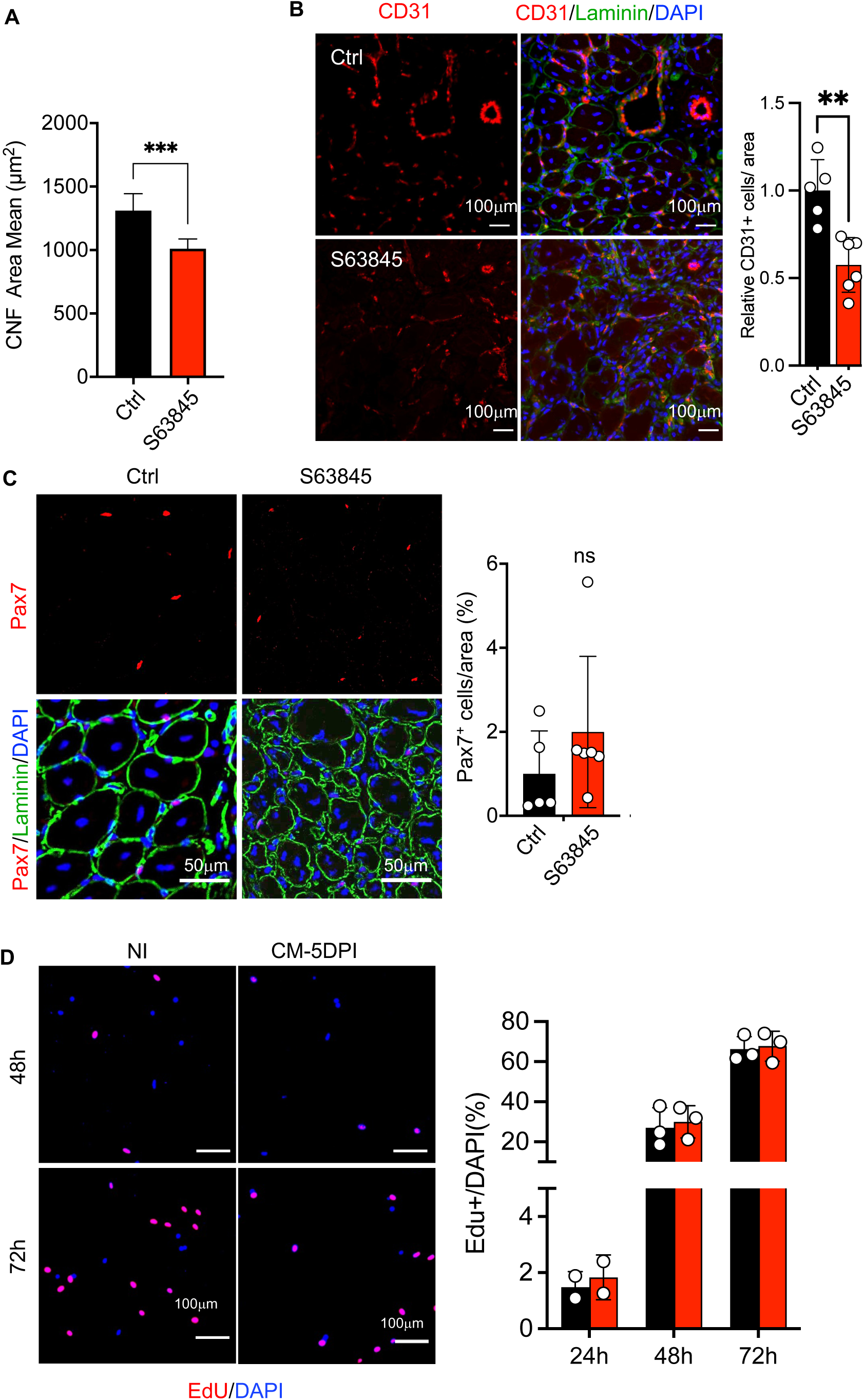
S63845 delays muscle regeneration. **A.** Mean area of myofibers with centralized nuclei (CNF) in the control group and the S63845-treated group. **B.** Representative images and quantifications of CD31 staining in TA muscles from the control group and the S63845-treated group. Scale bar=100μm. n=5-7. **C.** Representative images and quantifications of Pax7 staining in TA muscles from the control group and the S63845-treated group. Scale bar=50μm. n=5-7. **D.** Representative images and quantification of Edu staining in myoblasts cultured in CMs derived from FAPs either of NI or CTX 5dpi. at 48 hours and 72 hours after seeding. Scale bar=100μm, quantification was based on 3 biological repeats, every value was average of 400 cells/sample. Statistical significance was determined using Mann-Whitney test. Error bars represent mean ± SD. **p<0.01, *** p<0.001

**Supplementary Table 1.** Antibodies used un this study.

| Antibody | Source | Dilution |
| --- | --- | --- |
|  |  | IF: immunofluorescence<br>IHC: Immunohistochemistry |
| BrdU | Abcam (ab6326) | IF (1:200) |
| Lamin B1 | Abcam (ab16048) | IF (1:2000) |
| MCL1 | Abcam (ab32087) | IF (1:100) |
| γH2AX | Millipore (05-636) | IF (1:200) |
| Mf20 | DSHB | IF (1:30) |
| eMyHC | DSHB (F1.652-s) | IF (1:50) |
| Cleaved Caspase-3 | Cell Signaling (9661) | IF (1:300) |
| Laminin | Sigma-Aldrich (L9393) | IF (1:500) |
| F4/80 | Fisher Scientific (MA516624) | IHC (1:100) |
| PDGFRα | R&D (AF162) | IF (1:200) |
| Pax7 | DSHB | IF (1:20) |
| CD31 | Abcam (ab7388) | IF (1:100) |
| Donkey anti-mouse DyLight 488 | ImmunoReagents (DkxMu-003-D488NHSX) | IF (1:500) |
| Alexa Fluor 555 Goat anti-mouse | Invitrogen (A21124) | IF (1:1000) |
| Alexa Fluor 555 Goat anti-rabbit | Invitrogen (A21428) | IF (1:1000) |
| Alexa Fluor 633 Goat anti-Rabbit | Invitrogen (A21070) | IF (1:1000) |

**Supplementary Table 2.**
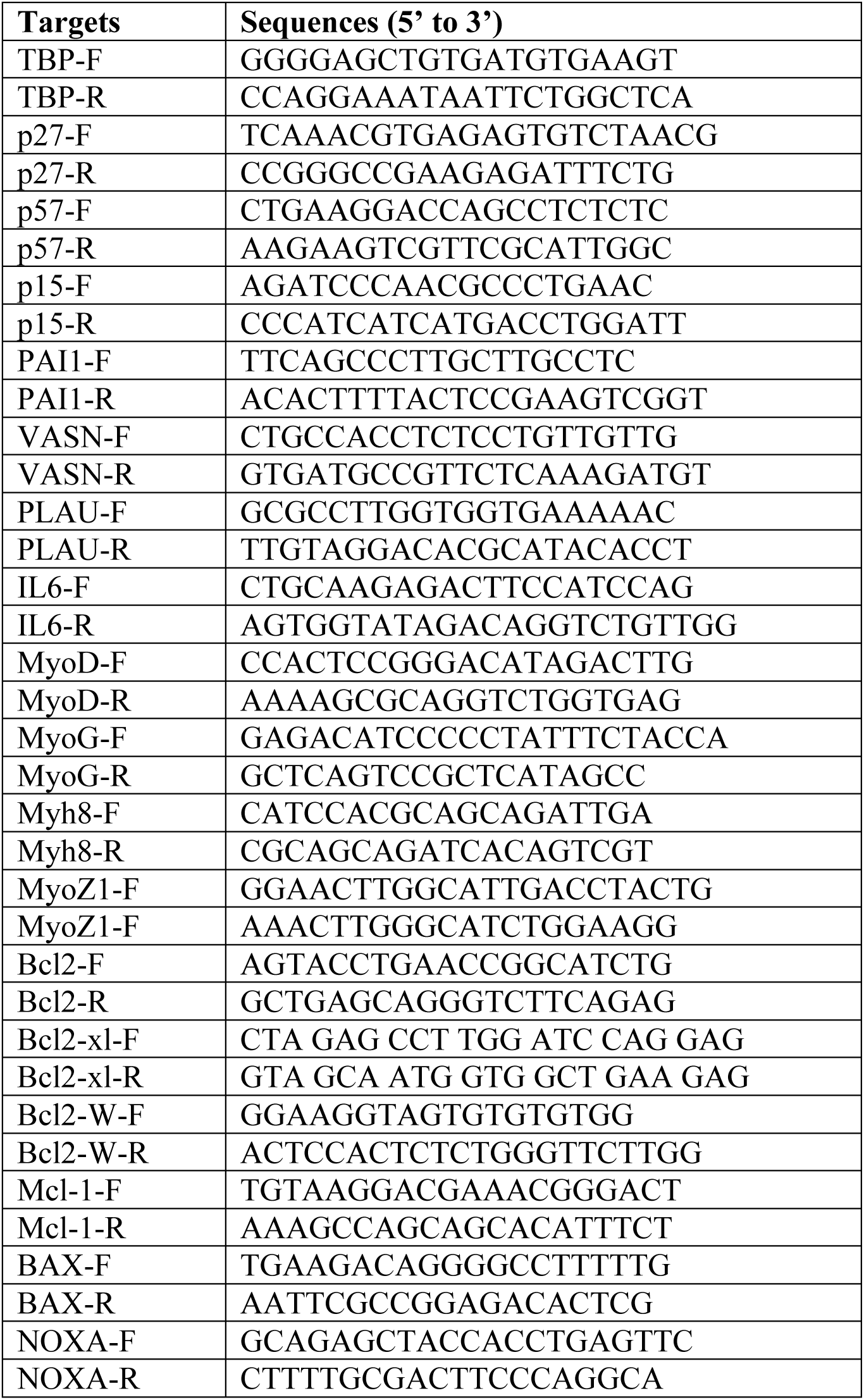
Primers for qPCR in this study.

## REFERENCE

1 Gorgoulis, V. et al. Cellular Senescence: Defining a Path Forward. Cell 179, 813–827, doi:10.1016/j.cell.2019.10.005 (2019).

2 Birch, J. & Gil, J. Senescence and the SASP: many therapeutic avenues. Genes Dev 34, 1565–1576, doi:10.1101/gad.343129.120 (2020).

3 Kirschner, K., Rattanavirotkul, N., Quince, M. F. & Chandra, T. Functional heterogeneity in senescence. Biochem Soc Trans 48, 765–773, doi:10.1042/BST20190109 (2020).

4 Munoz-Espin, D. et al. Programmed cell senescence during mammalian embryonic development. Cell 155, 1104–1118, doi:10.1016/j.cell.2013.10.019 (2013).

5 Storer, M. et al. Senescence is a developmental mechanism that contributes to embryonic growth and patterning. Cell 155, 1119–1130, doi:10.1016/j.cell.2013.10.041 (2013).

6 Demaria, M. et al. An essential role for senescent cells in optimal wound healing through secretion of PDGF-AA. Dev Cell 31, 722–733, doi:10.1016/j.devcel.2014.11.012 (2014).

7 Borghesan, M., Hoogaars, W. M. H., Varela-Eirin, M., Talma, N. & Demaria, M. A Senescence-Centric View of Aging: Implications for Longevity and Disease. Trends Cell Biol 30, 777–791, doi:10.1016/j.tcb.2020.07.002 (2020).

8 Evano, B. & Tajbakhsh, S. Skeletal muscle stem cells in comfort and stress. NPJ Regen Med 3, 24, doi:10.1038/s41536-018-0062-3 (2018).

9 Sousa-Victor, P. et al. Geriatric muscle stem cells switch reversible quiescence into senescence. Nature 506, 316-321, doi:10.1038/nature13013 (2014).

10 Garcia-Prat, L. et al. Autophagy maintains stemness by preventing senescence. Nature 529, 37–42, doi:10.1038/nature16187 (2016).

11 Moiseeva, V. et al. Senescence atlas reveals an aged-like inflamed niche that blunts muscle regeneration. Nature 613, 169–178, doi:10.1038/s41586-022-05535-x (2023).

12 Dungan, C. M. et al. Deletion of SA beta-Gal+ cells using senolytics improves muscle regeneration in old mice. Aging Cell 21, e13528, doi:10.1111/acel.13528 (2022).

13 Zhang, X. et al. Characterization of cellular senescence in aging skeletal muscle. Nat Aging 2, 601–615, doi:10.1038/s43587-022-00250-8 (2022).

14 Le Roux, I., Konge, J., Le Cam, L., Flamant, P. & Tajbakhsh, S. Numb is required to prevent p53-dependent senescence following skeletal muscle injury. Nat Commun 6, 8528, doi:10.1038/ncomms9528 (2015).

15 Chiche, A. et al. Injury-Induced Senescence Enables In Vivo Reprogramming in Skeletal Muscle. Cell Stem Cell 20, 407–414 e404, doi:10.1016/j.stem.2016.11.020 (2017).

16 von Joest, M. et al. Amphiregulin mediates non-cell-autonomous effect of senescence on reprogramming. Cell Rep 40, 111074, doi:10.1016/j.celrep.2022.111074 (2022).

17 Saito, Y., Chikenji, T. S., Matsumura, T., Nakano, M. & Fujimiya, M. Exercise enhances skeletal muscle regeneration by promoting senescence in fibro-adipogenic progenitors. Nat Commun 11, 889, doi:10.1038/s41467-020-14734-x (2020).

18 Young, L. V. et al. Muscle injury induces a transient senescence-like state that is required for myofiber growth during muscle regeneration. FASEB J 36, e22587, doi:10.1096/fj.202200289RR (2022).

19 Debacq-Chainiaux, F., Erusalimsky, J. D., Campisi, J. & Toussaint, O. Protocols to detect senescence-associated beta-galactosidase (SA-betagal) activity, a biomarker of senescent cells in culture and in vivo. Nat Protoc 4, 1798–1806, doi:10.1038/nprot.2009.191 (2009).

20 Behmoaras, J. & Gil, J. Similarities and interplay between senescent cells and macrophages. J Cell Biol 220, doi:10.1083/jcb.202010162 (2021).

21 Freund, A., Laberge, R. M., Demaria, M. & Campisi, J. Lamin B1 loss is a senescence-associated biomarker. Mol Biol Cell 23, 2066–2075, doi:10.1091/mbc.E11-10-0884 (2012).

22 Hall, B. M. et al. p16(Ink4a) and senescence-associated beta-galactosidase can be induced in macrophages as part of a reversible response to physiological stimuli. Aging (Albany NY) 9, 1867–1884, doi:10.18632/aging.101268 (2017).

23 Takeuchi, S. et al. Intrinsic cooperation between p16INK4a and p21Waf1/Cip1 in the onset of cellular senescence and tumor suppression in vivo. Cancer Res 70, 9381–9390, doi:10.1158/0008-5472.CAN-10-0801 (2010).

24 Chinzei, N. et al. P21 deficiency delays regeneration of skeletal muscular tissue. PLoS One 10, e0125765, doi:10.1371/journal.pone.0125765 (2015).

25 Saul, D. et al. A new gene set identifies senescent cells and predicts senescence-associated pathways across tissues. Nat Commun 13, 4827, doi:10.1038/s41467-022-32552-1 (2022).

26 Giordani, L. et al. High-Dimensional Single-Cell Cartography Reveals Novel Skeletal Muscle-Resident Cell Populations. Mol Cell 74, 609–621 e606, doi:10.1016/j.molcel.2019.02.026 (2019).

27 Hamilton, T. G., Klinghoffer, R. A., Corrin, P. D. & Soriano, P. Evolutionary divergence of platelet-derived growth factor alpha receptor signaling mechanisms. Mol Cell Biol 23, 4013–4025, doi:10.1128/MCB.23.11.4013-4025.2003 (2003).

28 Lemos, D. R. et al. Nilotinib reduces muscle fibrosis in chronic muscle injury by promoting TNF-mediated apoptosis of fibro/adipogenic progenitors. Nat Med 21, 786–794, doi:10.1038/nm.3869 (2015).

29 Chang, J. et al. Clearance of senescent cells by ABT263 rejuvenates aged hematopoietic stem cells in mice. Nat Med 22, 78–83, doi:10.1038/nm.4010 (2016).

30 Troiani, M. et al. Single-cell transcriptomics identifies Mcl-1 as a target for senolytic therapy in cancer. Nat Commun 13, 2177, doi:10.1038/s41467-022-29824-1 (2022).

31 Kohli, J. et al. Targeting anti-apoptotic pathways eliminates senescent melanocytes and leads to nevi regression. Nat Commun 13, 7923, doi:10.1038/s41467-022-35657-9 (2022).

32 Joe, A. W. et al. Muscle injury activates resident fibro/adipogenic progenitors that facilitate myogenesis. Nat Cell Biol 12, 153–163, doi:10.1038/ncb2015 (2010).

33 Lukjanenko, L. et al. Aging Disrupts Muscle Stem Cell Function by Impairing Matricellular WISP1 Secretion from Fibro-Adipogenic Progenitors. Cell Stem Cell 24, 433–446 e437, doi:10.1016/j.stem.2018.12.014 (2019).

34 Prieto, L. I. et al. Senescent alveolar macrophages promote early-stage lung tumorigenesis. Cancer Cell 41, 1261–1275 e1266, doi:10.1016/j.ccell.2023.05.006 (2023).

35 Haston, S. et al. Clearance of senescent macrophages ameliorates tumorigenesis in KRAS-driven lung cancer. Cancer Cell 41, 1242–1260 e1246, doi:10.1016/j.ccell.2023.05.004 (2023).

36 Cohn, R. L., Gasek, N. S., Kuchel, G. A. & Xu, M. The heterogeneity of cellular senescence: insights at the single-cell level. Trends Cell Biol 33, 9–17, doi:10.1016/j.tcb.2022.04.011 (2023).

## REFERENCE

1. Serrano, M. et al. Role of the INK4a Locus in Tumor Suppression and Cell Mortality. Cell 85, 27– 37 (1996).

2. Mukherjee, N. et al. MCL1 inhibitors S63845/MIK665 plus Navitoclax synergistically kill difficult-to-treat melanoma cells. Cell Death Dis 11, 443 (2020).

3. Muscle Stem Cells: Methods and Protocols. vol. 1556 (Springer New York, 2017).

4. Liu, L., Cheung, T. H., Charville, G. W. & Rando, T. A. Isolation of skeletal muscle stem cells by fluorescence-activated cell sorting. Nat Protoc 10, 1612–1624 (2015).

5. Chiche, A. et al. Injury-Induced Senescence Enables In Vivo Reprogramming in Skeletal Muscle. Cell Stem Cell 20, 407-414.e4 (2017).

6. von Joest, M. et al. Amphiregulin mediates non-cell-autonomous effect of senescence on reprogramming. Cell Rep 40, 111074 (2022).

7. Korsunsky, I. et al. Fast, sensitive and accurate integration of single-cell data with Harmony. Nat Methods 16, 1289–1296 (2019).

8. Cazin, C., Chiche, A. & Li, H. Evaluation of Injury-induced Senescence and In Vivo Reprogramming in the Skeletal Muscle. JoVE 56201 (2017) doi:10.3791/56201.

9. Chantrel, J., Chen, C., Zhang, J. & Li, H. Protocol for quantitative evaluation of the impact of paracrine senescence on cellular reprogramming in cultured cells and mouse models. STAR Protoc 4, 102106 (2023).

10. Mayeuf-Louchart, A. et al. MuscleJ: a high-content analysis method to study skeletal muscle with a new Fiji tool. Skeletal Muscle 8, 25 (2018).

